# Immediately following the onset of active whisking, the input layer of barrel cortex exhibits a transient period of increased excitability that depends on prior experience

**DOI:** 10.1101/2024.06.04.597353

**Authors:** Molly C. Shallow, Lucy Tian, Bryan T. Higashikubo, Hudson Lin, Katheryn B Lefton, Siyu Chen, Joseph D. Dougherty, Joe P. Culver, Mary E. Lambo, Keith B. Hengen

## Abstract

The development of motor control over sensory organs is a critical milestone, enabling active exploration and shaping of the sensory environment. Whether the onset of sensory organ motor control directly influences the development of corresponding sensory cortices remains unknown. Here, we confirm and exploit the late onset of whisking behavior in mice to address this question in the somatosensory system. Using ex vivo electrophysiology, we describe a transient increase in the intrinsic excitability of excitatory neurons in layer IV of the barrel cortex, which processes whisker input, immediately following the onset of active whisking around on postnatal days 13 and 14. This increase in neuronal gain is specific to layer IV, independent of changes in synaptic strength, and requires prior sensory experience. Further, these effects are not expressed in inhibitory interneurons in barrel cortex. The transient increase in excitability is not evident in layer II/III of barrel cortex or in the visual cortex upon eye opening, suggesting a unique interaction between the development of active sensing and the thalamocortical input layer in the somatosensory isocortex. Predictive modeling indicates that, immediately following the onset of active whisking, changes in active membrane conductances alone can reliably distinguish neurons in control but not whisker-deprived hemispheres. Our findings demonstrate an experience-dependent, lamina-specific refinement of neuronal excitability tightly linked to the emergence of active whisking. This transient increase in the gain of the thalamic input layer coincides with a critical period for synaptic plasticity in downstream layers, suggesting a role in cortical maturation and sensory processing.

## Introduction

Isocortical microcircuits give rise to conscious perception^1–3^, adaptable interaction with stimuli^4–7^, and rapidly adjust to the environment into which an individual is born^8–12^. As a result, understanding the combination of internal processes and external influences that contribute to the development of isocortical networks is a key question with broad consequences, both neurobiological^13^ and societal^14,15^ in form. However, it is challenging to identify and control the features of the external world that drive specific isocortical circuits. As a result, it can be difficult to pinpoint the mechanisms by which isocortical development interacts with the environment. With this in mind, the mouse whisker-barrel system is an ideal model for studying isocortical postnatal development, specifically the interaction of experience with internal programs. The whisker-barrel system is characterized by the emergence of topographically organized maps whose architecture is notably accessible for two reasons; *first*, under normal conditions, the stereotyped rows and columns of whisker pads on the face are distinctly visible in the cellular architecture in layer IV (i.e., “barrels” in S1^16^), and *second*, it is easy to manipulate highly specific inputs by simply removing the desired subset of whiskers — each of which map onto a single barrel^17^. Because of this accessibility, the whisker-barrel system has been utilized to uncover broad principles of development, plasticity, sensory coding, and isocortical microcircuitry^18–23^.

While the structural development of barrels in the first days of life has been extensively studied^24,25^, less is known about the isocortical impact of the onset of active whisking around postnatal day 14 (P14)^26^. The active manipulation of sensory organs to explore the world (such as sniffing with the nose), is a prominent feature of most sensory systems^27–30^. This is true of tactile information; from the first moments of life, cutaneous somatosensation is driven by limb movement and locomotion^31^. In contrast to the rest of the body, however, an animal’s whiskers are not actively controlled during the first weeks of life^32–34^. While the barrels are formed and innervated prior to and shortly after birth^18,20,32,35,36^, passive input through the whiskers in the first ∼two weeks establishes connectivity from the barrels (layer IV) to layers II/III^37^. A few days before the onset of active whisking, there is a brief period of “symmetric whisking”, signaling an impending change from passive to active sensing^32^. Then, analogous to the binary shift that comprises eye opening, around P14, it is believed that mice commence control of individual whiskers for exploration of objects and the environment^32^. Perhaps unsurprisingly, the emergence of complex whisking behavior coincides with significant Hebbian modification in layers II/III in and after the third week of life33,38,39.

Despite phenomenological parallels with eye opening, the onset of active whisking differs in at least one crucial aspect: by P14, the critical period for barrel formation and structural plasticity is largely understood to be closed^24,35,40^. In contrast, the critical period for ocular dominance plasticity begins ∼10 d after eye opening^41^. This delay between the maturation of sensory transduction and the behavioral control of whiskers provides a unique opportunity to isolate the role of motor control in sensory system development. In other words, the barrel cortex around P14 is distinctive— it presents a structurally mature circuit exposed to a developmentally-timed onset of organized input^16,42^. The default possibility is that the commencement of active whisking ( ∼P14) simply introduces complex stimuli into established circuitry. In other words, the system is assembled prior to active, complex input. In this case, motor control of sensory organs indirectly refines sensory cortex by determining stimuli — largely reflected by plasticity in layers II/III^33,38,39^. In this case, the prediction is that neurophysiological properties of barrel cortex, particularly in input layers, are fixed by the time of active whisking^24,35,40^. Alternatively, sensory cortices could cooperate with key events in motor development. In this case, the onset of whisker movement should accompany a parallel change in barrel fields that is positioned to facilitate information processing. Whether the barrel cortex actively cooperates with the onset of whisking behavior and consequent input is unknown^43^. Support for this hypothesis would comprise an experience-dependent alteration in barrel cortex roughly coincident with the emergence of active whisking.

Based on prior work, it is reasonable to suggest that a plastic change in layer IV barrels is unlikely on or about P14. Barrel structure and synaptic connectivity between layers are established earlier in life^40^, and the relative immutability of sensory cortices after critical period closure is a general phenomenon^44^. While plasticity isn’t limited to the synapse — intrinsic plasticity is regularly observed in many systems — non-synaptic changes almost always accompany synaptic plasticity^45–49^. In this context, we sought to answer a simple question: is there evidence of a plastic interaction between layer IV barrel cortex and the onset of whisker motor control?

To address this, measured the ontogeny of whisking behavior in mice, and employed ex vivo slice recordings in layers II/III and IV of barrel cortex as well as in layer IV in primary visual cortex. We measured intrinsic excitability and synaptic strength throughout this key developmental time period to assess the possibility of plastic change in the barrels at the onset of active whisking. We identify a robust, non-synaptic form of experience-dependent plasticity that occurs immediately following the onset of active whisking. This transient event is specific to excitatory neurons in layer IV barrel cortex.

## Results

To explore the possibility that neurons in the barrel fields of layer IV S1 (S1_BF_) show evidence of a plastic change around the onset of active whisking (approximately postnatal days 13-14; P13-14), we made acute, ex vivo brain slices (350 *µ*m) containing layer IV barrels from mice (N = 82 animals; Fig. 2A,B). We conducted whole-cell patch-clamp recordings from excitatory spiny stellate neurons, identified by morphological characteristics and confirmed by post hoc imaging (^50,51^, Fig. 2B). To capture possible changes around the onset of active whisking, slices were made intermittently between P11 and P19 (Fig. 2C). We assessed intrinsic excitability via standard F/I curves (firing rate per current step;^49^), and we estimated the distribution of post-synaptic strengths by measuring miniature excitatory post-synaptic potentials (mEPSCs;^49,52^). In addition to stellate neurons in the barrels, we recorded excitatory pyramidal neurons in layers II/III of S1 (dorsal to the barrels;^53^), as well as two populations of NKX2.1-positive inhibitory interneurons—somatostatin (SST) and parvalbumin (PV) subtypes^54,55^.

### Developmental Onset of Active Whisking

We first sought to empirically determine the developmental timecourse of the emergence of active whisking behavior in C57BL6/J mice. This is necessary, as prior descriptions of a rapid transition around P14 are limited^32^. Phylogenetic differences aside, limited results in rats suggest an abrupt transition at P14^56^, while other rat work describes a gradual transition that is completed by ∼ P14^57^.

To characterize the timing and abruptness of active whisking onset in mice, we systematically analyzed whisking behavior from P7 to P16 (*n* = 10–28 animals per age). Animals were scored by an experimenter naive to prior evidence of developmental trajectories — note that, due to the dramatic body morphological changes that occur between P7 and P16, it is impossible to blind an observer to animal age. Animals were categorized as exhibiting either non-whisking, twitching, or active whisking behavior (Fig. 1A). Active whisking was observed first on P12, and was evident in ≥ 90% of animals at P14 (FIg. 1B). Binary logistic regression analysis demonstrated that age was a highly significant predictor of active whisking behavior (*β* = 2.54, *p* = 6.28 × 10*^−^*^8^, pseudo-*R*^2^ = 0.77) according to the equation 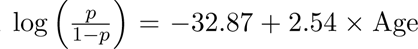 (Fig. 1C). This model predicts that the probability of active whisking reaches 50% at postnatal day 12.93 (95% CI: [12.69, 13.18]). Multinomial logistic regression analysis revealed distinct developmental trajectories for all three behavioral states (*χ*^2^ = 323.9, *df* = 2, *p* = 4.58 × 10^−71^, pseudo-*R*^2^ = 0.73). The transition to twitching behavior was modeled by the equation 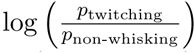= −19.69+2.01×Age (*p <* 0.001), while the transition to active whisking followed 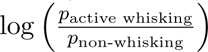 = −52.38+4.54×Age (*p <* 0.001).

**Figure 1:**
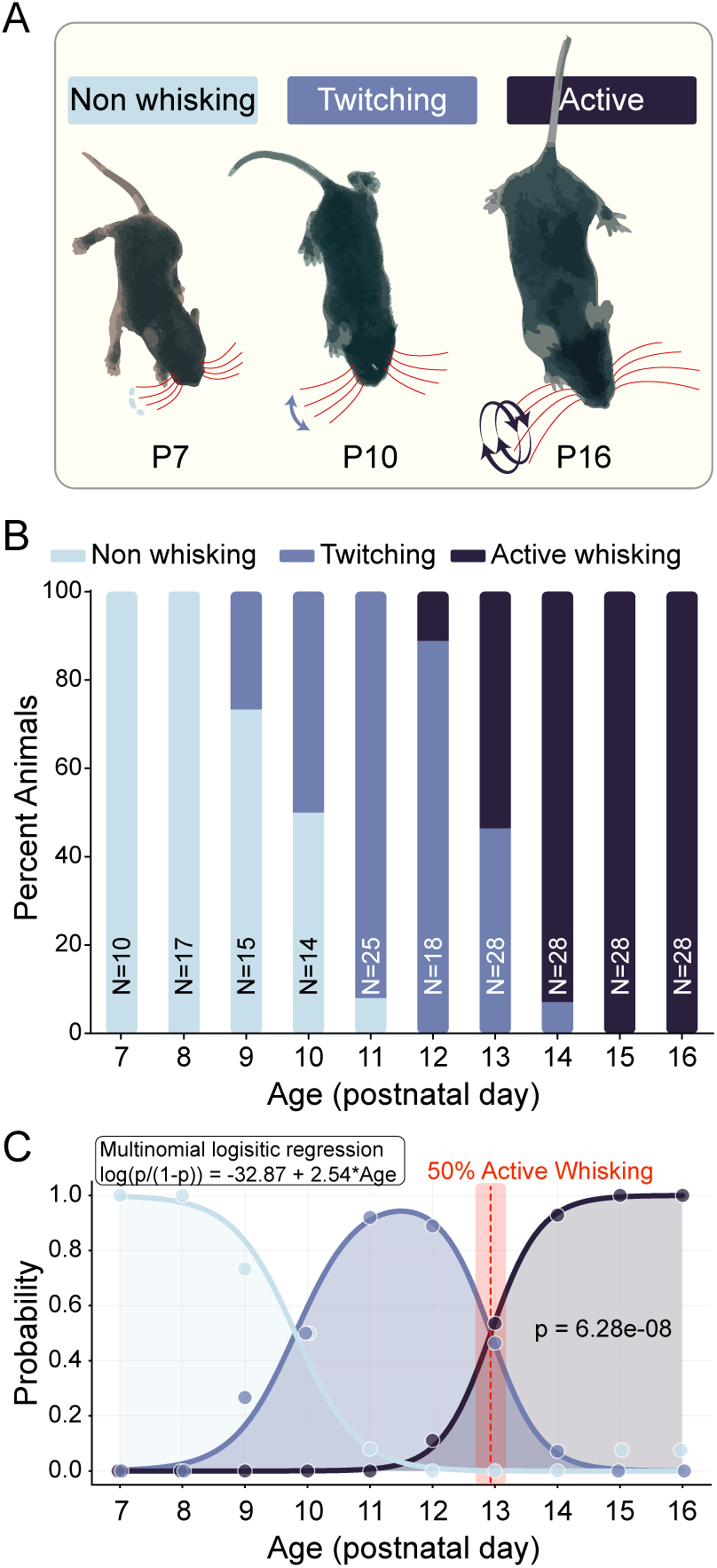
Developmental onset of active whisking behavior in mice. **A)** Representative images illustrating three categories of whisking behaviors. Red lines indicate whisker position, with blue arrows showing movement patterns. At P7, mice exhibit non-whisking behavior characterized by largely immobile whiskers. By P10, twitching behavior emerges, characterized by small, non-rhythmic whisker movements. At P16, mice display active whisking with large-amplitude, rhythmic, bilateral whisker movements. **B)** Stacked bar graph showing the percentage of animals exhibiting each whisking behavior across postnatal days P7– P16. Sample sizes are indicated for each age group. Non-whisking behavior (light blue) dominates at early ages (P7–P8), transitions to predominantly twitching behavior (medium blue) during intermediate ages (P9–P12), and shifts to active whisking (dark blue) at later ages (P13–P16). Complete transition to active whisking occurs by P15. **C)** Multinomial logistic regression analysis of the developmental transition. Light blue curve represents the probability of non-whisking behavior, medium blue curve represents twitching, and dark blue curve represents active whisking as a function of age. The binary logistic regression equation for active whisking probability is shown (log(*p/*(1 − *p*)) = −32.87 + 2.54 × Age, *p* = 6.28 × 10*^−^*^8^). The vertical red dashed line indicates the age at which active whisking probability reaches 50% (P12.93), with pink shading representing the 95% confidence interval. Gray shading indicates the age range where active whisking becomes the predominant behavior.

The coefficient for active whisking (4.54) was more than twice as large as the coefficient for twitching (2.01), indicating a substantially more abrupt developmental transition to active whisking. This is consistent with prior evidence in mice^32^. We observed near-complete expression (*>* 90%) of active whisking by P14 and total expression (100%) by P15. A warning of quasi-separation in the logistic model further supports the abrupt nature of this transition, with complete separation of behaviors at the upper age range. Together, these results provide strong statistical evidence that the emergence of active whisking follows a non-linear developmental trajectory with a rapid transition between P12 and P14. At least in C57BL6/J mice, active whisking appears to emerge through an abrupt developmental switch rather than through gradual onset. Crucially, across all examined animals, P14 thus represent the first day during which the barrel fields of S1 are exposed to information derived from active exploration with the whiskers.

### Excitability of barrel field excitatory neurons is increased on P14 (but not P12 or P16)

A fundamental property determining whether and how a neuron converts inputs into spikes is the intrinsic excitability of its membrane. A defining feature of isocortical physiology in early postnatal life is a progressive reduction in neuronal intrinsic excitability. This typically unfolds over weeks, and is characterized by increasing thresholds for action potential generation and decreasing resting membrane potentials. As isocortical neurons grow and develop more complex dendritic trees and form more synaptic connections, their input resistance typically decreases, which reduces their overall excitability (Fig. S1)^58–60^. Membrane excitability is assayed by injecting an increasing series of positive current steps and counting the number of action potentials generated by each step. The slope of the resultant curve—the F/I curve— thus describes the intrinsic excitability of a neuron.

Given the progressive decrease in excitability across development, the establishment of active whisking by P14 might reasonably be expected to drive no meaningful change in intrinsic excitability beyond that which is already unfolding. Alternatively, increased sensory behavior might result in a compensatory decrease in intrinsic excitability, as isocortical sensory neurons homeostatically regulate their excitability to counterbalance significant changes in input^45^. To evaluate this, we patched spiny stellate neurons (excitatory) in visually identified barrels at P11, 12, 14, 16, 18, and 19. We delivered ten 500 ms currents between 0 and 180 pA in steps of 20 pA (Fig. 2D). For quantification, we fit a linear model to each neuron’s F/I curve between 20 and 100 pA (all cells were active and not saturated in that range) and extracted the model slope as an estimate of intrinsic excitability. There was a significant main effect of age (*p* = 1.26*e* − 05, mixed-effects analysis of variance: slope ∼ age + animal id, where animal ID is included as a fixed effect). Posthoc comparisons of P14 to all other ages yielded *p* values between 0.0000064 and 0.013 (Fig. 2E). Examination of passive properties—tau, membrane resistance, capacitance, and resting membrane potential—revealed either no change or a progressive decline with age, as expected. There were no significant alterations in any passive properties on P14 (Fig. S1A-D). These results suggest that excitatory neurons in the input layer of S1_BF_ support an unexpected, transient increase in intrinsic excitability immediately following the onset of active whisking.

**Figure 2:**
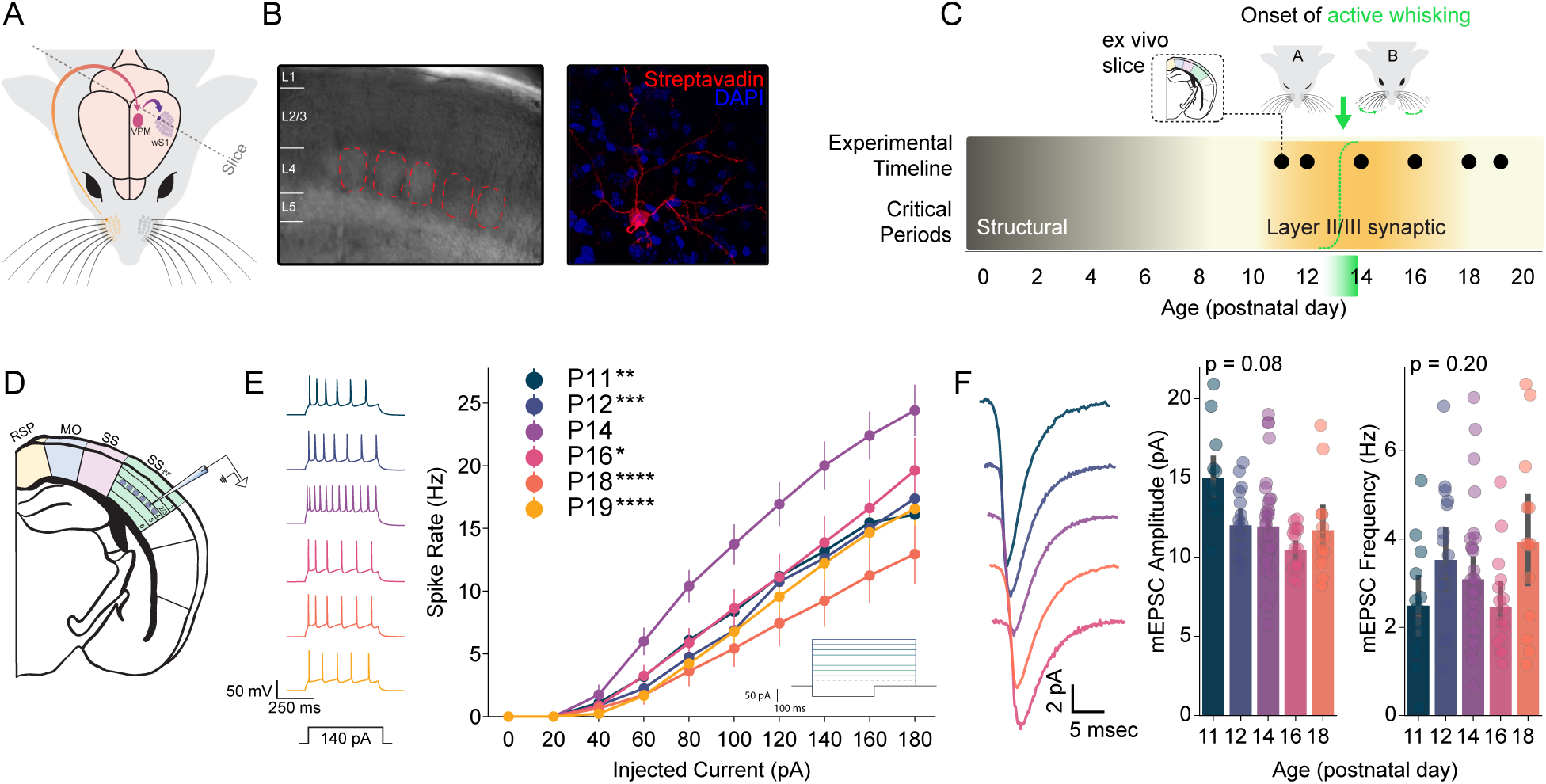
Around the onset of active whisking, spiny stellate neurons in layer 4 barrel cortex exhibit a transient increase in excitability. **A)** Experimental approach. Ex vivo slices were taken from the barrel field of primary somatosensory cortex (SS_BF_) from postnatal day 11 (P11) through P19. **B)** Layer IV barrels were visualized under differential interference contrast (DIC) imaging at 10x and individual excitatory cells were chosen under 40x magnification. After recording, cell morphology and location were visualized using streptavidin stains and confocal microscopy. **C)** Timeline of experiment. Outline of relevant critical periods for structural and synaptic plasticity as well as the timeline of recordings and onset of active whisking (dashed green sigmoid and fade). **D)** Location of intracellular patch recordings in SS_BF_. **E)** Immediately following the onset of active whisking (see Fig. 1), P14 F/I curves reveal a transient increase in the intrinsic excitability of Layer IV excitatory cells evident, right. Example traces from each age, left. **F)** mEPSC recorded from Layer IV excitatory cells do not show an accompanying synaptic strength change, right. Example mEPSC traces from each age, left. Printed p values are for the main effect of age. For F/I curves N=26 mice, N=108 cells. For mEPSCs N=22 mice, N=94 cells. \**p <* 0.05, ** *p <* 0.01, *** *p <* 0.001, ^****^ *p <* 0.0001

Intrinsic excitability plasticity is tightly linked with congruent changes in synaptic strength^45–49^. Thus, we reasoned that elevated excitability on P14 might indicate a generalized increase in synaptic strength — as would be expected of a modifiable circuit subjected to increased input. Alternatively, increased input might drive a compensatory decrease in synaptic strength^52,61–63^. To test these possibilities, we evaluated mEPSC amplitudes in S1_BF_ excitatory neurons between P11 and P18— note that mEPSC amplitude is reflective of both homeostatic and Hebbian plasticity^11,49,52,64^. Surprisingly, we found no evidence of any change in synaptic strength around P14 (Fig. 2F). Across the range of ages in question, mEPSC amplitude trended towards a decrease over time (*p* = 0.08; mixed-effects analysis of variance: mini amplitude ∼ age + animal id), which is consistent with postnatal isocortical maturation^65^. However, in contrast to intrinsic excitability, there was no change in mEPSC amplitude specific to P14 (P12 vs P14 *p* = 0.8915; P16 vs P14 *p* = 0.8515). mEPSC frequency, often taken as an indicator of presynaptic properties^66^, also showed no change over time (*p* = 0.20) and no specific change on P14 (P12 vs P14 *p* = 0.9934; P16 vs P14 *p* = 0.8216).

In addition to mEPSC amplitude, glutamatergic plasticity can alter subunit composition^67^. To evaluate this possibility, we examined the mEPSC waveform, as channel composition specifics determine waveform shape^68,69^. While AMPA-R subunit composition is expected to change across postnatal development^70^, and thus influence mEPSC waveform shape, we sought to test whether the kinetics of glutamate-mediated synaptic transmission change drastically around the onset of active whisking. We trained a classifier to predict age based on mEPSC waveform — should there be distinct, consistent differences in any aspect of the mEPSC waveform on P14, a classifier would learn this in a data-driven manner. We computed the mean mEPSC waveform for each neuron recorded from P11 to P18 and extracted the first four principal components, which encompassed the majority of the variance in the data (82.5%), allowing us to describe waveform characteristics without substantial information loss (Fig. S1E,F). We then trained a class-balanced logistic regression model to predict the age group based on the principal component scores (Stratified K-Fold cross-validation with 5 splits and 200 repeats). Between P11 and P18, the overall balanced accuracy was 48% (chance = 20%), an effect driven by classifier performance on P11 (62.7%) and P16 (77.5%) (Fig. S1G), consistent with a gradual, time-dependent shift in GluA subunit composition across development. However, on P14, performance was below chance (18.8%), suggesting that there is no discernible shift in the dynamics of glutamatergic synaptic transmission driven by or in anticipation of onset of active whisking. Examination of PC1 loadings indicates that the differentiability of ages (i.e., P11 and P18) was largely driven by changes in the falling phase of the mEPSC (Fig. S1H).

Taken together, these data suggest that, around the onset of active whisking, there is a plastic event in layer IV barrel cortical excitatory neurons. Surprisingly, synaptic strength and kinetics are unaffected. This raises the possibility of a brief, developmentally-timed increase in the gain of input from VPM thalamus to isocortex that cooperates with the initiation of a related motor program. However, it is important to assess the cell-type specificity of this phenomenon, as an equivalent change in inhibitory neurons in layer IV S1_BF_ might mitigate increased excitatory gain. In other words, a generalized, network-wide increase in excitability could be self-stabilizing.

### Barrel inhibitory neurons are unaffected by the onset of active whisking

To investigate the possibility that increased intrinsic excitability is a general feature of barrel cortex around P14, we recorded F/I curves from parvalbumin-positive inhibitory interneurons (PV cells), which represent the most common inhibitory subclass in isocortex^61,71–74^. However, parvalbumin protein begins expression at approximately P14^75^, making it challenging to identify PV interneurons in younger animals. To circumvent this, we performed a genetic cross between two transgenic mouse lines. The first line, C57BL/6J-Tg(Nkx2-1-cre)2Sand/J^76^, expresses Cre recombinase under the control of the Nkx2-1 promoter, which targets forebrain GABAergic interneurons. The second line, TRAP (EGFP/RPL10A^77^), expresses EGFP in the presence of Cre recombinase. This combination results in EGFP expression in Nkx2.1 positive cells. Crucially, Nkx2.1 is a transcription factor that participates in the differentiation of PV and somatostatin (SST) interneurons^54,55^ (Fig. 3A). Within the Nkx2.1-positive population, PV and SST interneurons can be differentiated by assessing the presence/absence of SST immunoreactivity (Fig. 3B) and by electrophysiological properties (Fig. 3C)^78^.

**Figure 3:**
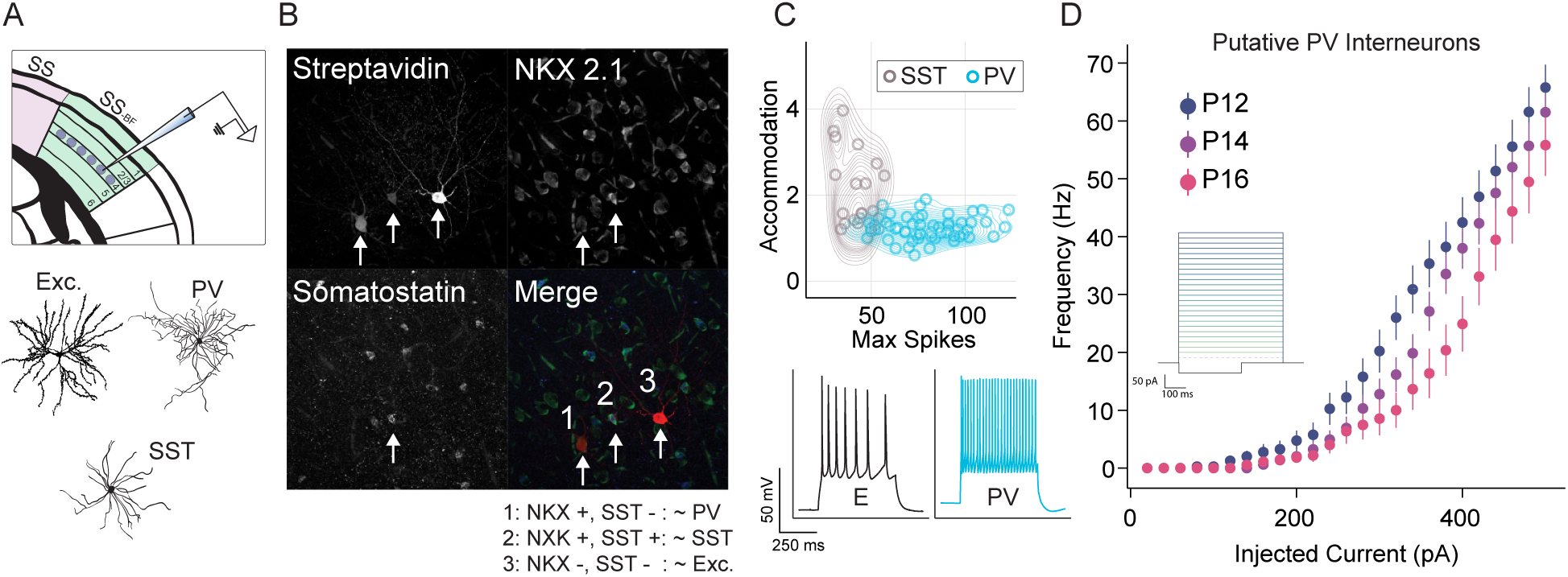
Increased excitability at the onset of active whisking is specific to excitatory neurons. **A)** Experimental approach. To fluorescently label a subset of isocortical inhibitory interneurons, Nkx2.1-Cre mice were crossed with TRAP mice, which express EGFP-tagged ribosomal protein L10a under a Cre-dependent promoter. (Top) In this context, Nkx2.1 positive neurons in Layer IV barrels were recorded on postnatal day 12 (p12) through p16. (Bottom) Subsequent staining for somatostatin (SST) protein allowed for the separation of three cell types in our dataset: excitatory, parvalbumin (NKX positive, SST negative), and SST (NKX and SST positive). **B)** Example triple label image demonstrating the separation of the three putative cell types. Streptavidin is loaded in the recording electrode and thus labels cells that were patched (intensity reflects the duration of patch). Nkx2.1 labels PV and SST interneurons in the isocortex. Somatostatin label is used to define SST interneurons. Merge demonstrates the molecular logic of putative cell type identification. **C)** PV and SST subpopulations of Nkx2.1 positive neurons are differentiable based on electrophysiological properties, accommodation and the peak instantaneous firing rate. (Top) Kernel density estimate and scatter of Nkx2.1 positive neurons’ accommodation and max firing, colored by PV/SST classification. (Bottom) Example traces from an excitatory neuron (E) and putative PV interneuron in response to a 160 pA current step. **D)** F/I curve of putative PV interneurons in Layer IV barrel cortex as a function of age reveals no increase in intrinsic excitability on p14. For F/I curves, N=15 mice, N=40 cells.

We recorded F/I curves in Nkx2.1-positive inhibitory interneurons on P12, P14, and P16 in a total of 15 mice. In comparison to excitatory neurons, Nkx2.1-positive neurons had higher spike thresholds and continued to increase their activity at current injections above 200 pA without depolarization blockade. Thus, inhibitory F/I curves were assessed from 0 and 500 pA in steps of 20 pA (Fig. 3D inset). We divided putative PV and putative SST subpopulations based on accommodation (a progressive reduction in firing during a sustained stimulus) and firing rate per current injection. In stark contrast to excitatory neurons, putative PV interneurons showed no sign of increased excitability on P14 (3D), and there were no signs of specific changes in passive properties on P14 (Fig. S2). These data suggest that the plasticity observed in excitatory neurons is not only distinct in its singular regulation of membrane excitability, but is also cell-type specific.

### A subset of electrophysiological features contribute to intrinsic plasticity at P14

Mechanisms controlling intrinsic excitability fall into three categories. First, passive conductances comprise ion channels that are constantly open (at rest) and allow ions to flow based on their electrochemical gradients. These channels maintain the resting membrane potential and contribute to the overall resistance and capacitance of the membrane, affecting how the cell responds to incoming stimuli without directly participating in the generation of action potentials. Second, active conductances comprise ion channels that open and close in response to changes in membrane potential. These conductances actively regulate the flow of ions across the membrane. They include voltage-gated channels like Na^+^, K^+^, and Ca^2+^ channels, which generate dynamic responses to depolarization. Third, spiking properties define the output signals of neurons and include action potential threshold, the frequency and pattern of firing, and the adaptability of response to sustained inputs.

To identify candidate mechanisms that might account for the increase in intrinsic excitability in barrel excitatory neurons on P14, we extracted measures reflecting passive conductances (resting membrane potential, input resistance, membrane time constant—tau, and capacitance) (Fig. 4A-E), active conductances (dynamic input resistance and time to first spike) (Fig. 4F,G), and spiking features (after hyperpolarization, spike frequency adaptation, spike threshold, and instantaneous frequency)(Fig. 4H-J). We employed a predictive modeling approach to ask which, if any, of these three feature sets was sufficient to accurately classify the age of withheld neurons, with particular attention to performance on P14. In simple terms, a computational model attempted to learn to predict an animal’s age based only on a small set of electrophysiological features from a neuron. Trained on active conductances, logistic regression models accurately classified withheld P14 neurons at 76.86 ± 1.02% accuracy (chance = 16.7%) (Fig. 4K). Trained on passive properties, models accurately identified P14 neurons at 35.1.± 1.16%, and trained on spiking features, models performed at 40.9.± 1.2% on P14 (Fig. 4L,M). The highest performance across all three models was P14 active conductances. Passive properties were most effective at predicting P11 (52%), while spiking features’ effectiveness peaked on P16 at 56%. Undoubtedly, ion channel expression changes across development and as a feature of experience— assuming a stereotyped developmental timeline, it is unsurprising that a classifier can learn signatures to discriminate cells collected only 48 h apart. However, it is noteworthy that the only set of features that meaningfully differentiates neurons on P14 is the active conductances, suggesting that the underlying channels interact with the onset of whisker motor control.

**Figure 4:**
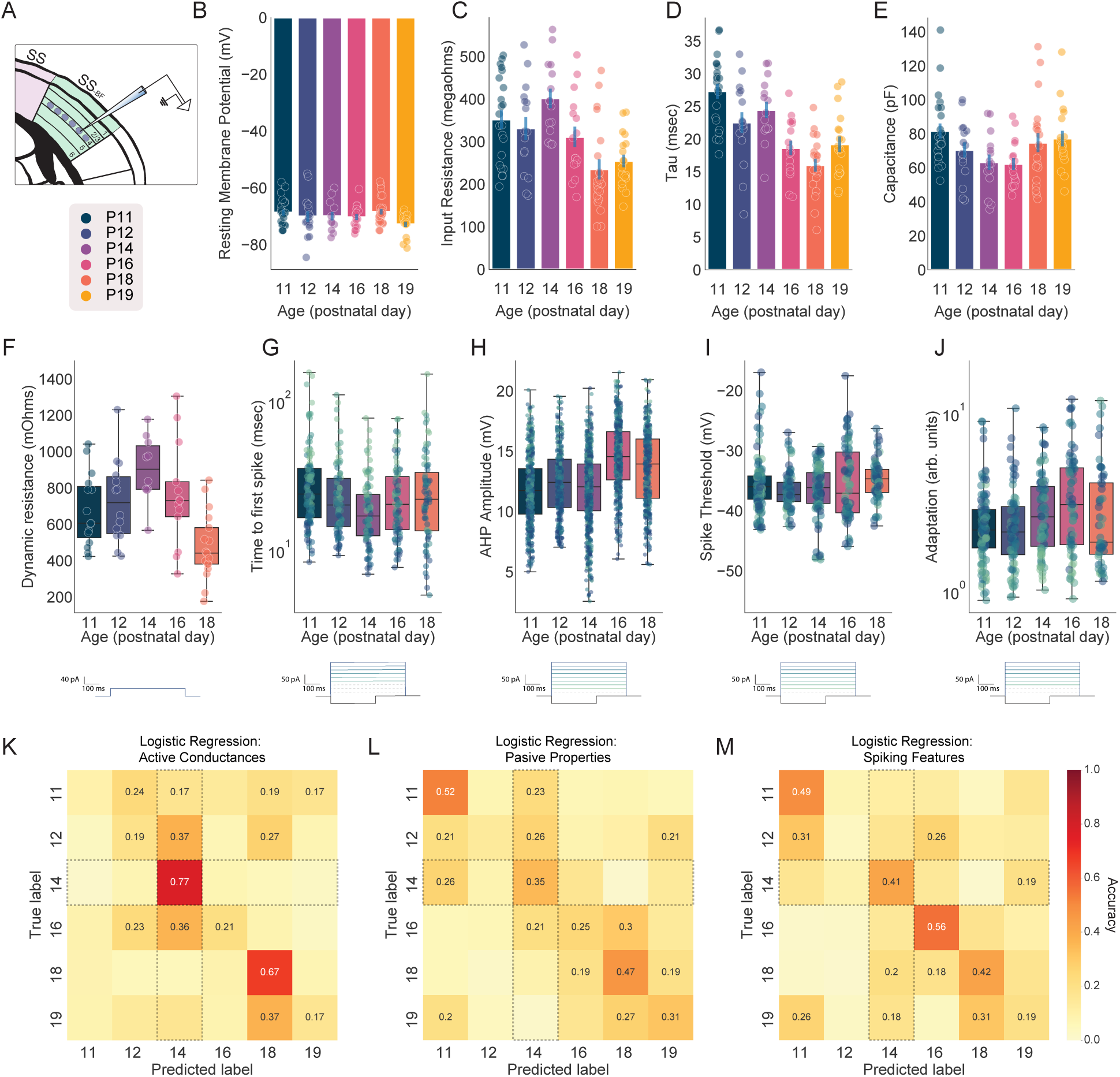
Plasticity in active conductances underlies increased excitability on P14. **A)** Illustra-tion of recording from excitatory neurons in Layer IV of the barrel fields of primary somatosensory cortex (SS_BF_). Legend indicates recorded ages and applies to the B-J. B-E) Passive properties of SS_BF_ L4 exci-tatory neurons as a function of age: **B)** resting membrane potential, **C)** input resistance (measured with a small hyperpolarizing current step), **D)** membrane time constant (tau), and **E)** membrane capacitance. **E)** Dynamic input resistance as a function of age. Note that this is a measure of the change in membrane resistance in response to a small positive current, in this case a 40 pA pulse. The voltage deflection is divided by the current (40 pA)— higher values suggest a less leaky membrane. **G)** Time to first spike as a function of age. **H)** Afterhyperpolarization (AHP) amplitude as a function of age. **I)** Spike threshold as a function of age. **J)** Spike frequency adaptation as a function of age— this refers to a decrease in firing rate over the course of a square wave. **K-M)** Logistic regression was used to predict animal age as a function of three features sets measured in each neuron: active conductances (dynamic input resistance and time to first spike), passive properties (membrane resistance, membrane potential, capacitance, tau), and spiking features (AHP, spike frequency adaptation, and instantaneous frequency). **K)** Confusion matrix— correctly classified neurons fall on the diagonal— for active conductances. Note that chance is 16.67% and that P14 is robustly identifiable. **L)** Same as K but for passive properties. **M)** Same as K but for spiking features.

### Region- and layer-specificity of intrinsic plasticity

Our findings thus far could represent a) a generalized, feed-forward Hebbian response to increased thamocortical input, b) a generalized feature of primary sensory cortical system upon the intro-duction of complex stimuli, or c) a process unique to the barrel fields. In the first case, one would expect that results similar to layer IV should be evident at the next layer of operation (i.e., layer II/III). In the second case, a parallel result should be observed in the primary visual cortex, as this also undergoes an abrupt transition upon eye opening. In the last case, our results should be evident only in layer IV barrel cortex.

We recorded F/I curves in excitatory pyramidal neurons in S1 layer II/III (immediately dorsal to the layer IV barrels) on P12, P14, and P16 (N = 11 mice). There was a significant main effect of age (*p* = 0.026; Fig. 5A), however this was explained by a decrease in excitability from P12 to P14 (*p* = 0.026) and P12 to P16 (*p* = 0.128). These data suggest that the effects observed in layer IV do not generalize to the rest of the barrel cortical circuit.

**Figure 5:**
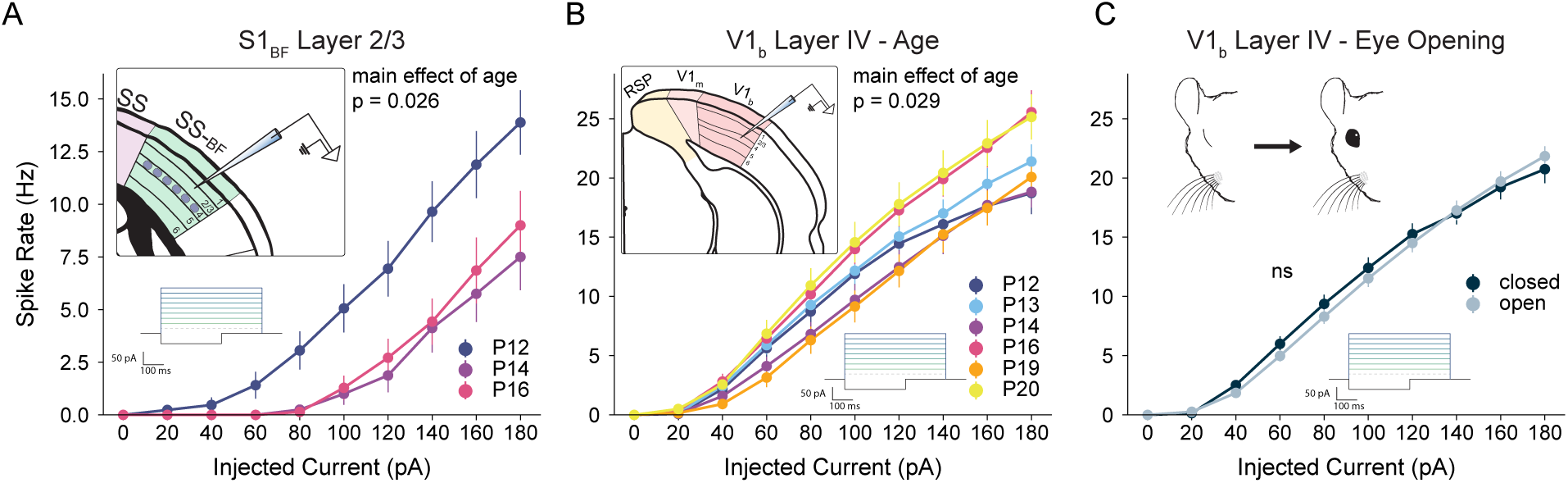
Increased excitability at P14 is not a general property of sensory cortices. **A)** Inset: Illustration of recording from excitatory neurons in Layer 2/3 of primary somatosensory cortex. Excitatory F/I curves for ages P12, P14, and P16. At P14, L2/3 neurons exhibit reduced excitability compared to P12. **B)** Excitatory F/I curve of excitatory L4 neurons in the binocular portion of primary visual cortex (V1_b_) for ages P12, P13, P14, P16, P19, and P20. At P14, neuronal excitability in V1_b_ is unremarkable. **C)** Mouse eye opening occurs between P10 and P14. F/I curve showing the data presented in B as a function of whether an animal’s eyes were open or closed. For Layer 2/3 F/I curves, N=11 mice, N=47 cells. For V1 F/I curves, N=19 mice, N=168 cells.

The visual system provides a similar opportunity to study the sudden introduction of complex stimuli into a cortical sensory network, as animals transition from a state of closed eyes to open in a relatively tight temporal window between P12 and P14^79^. To ask if the intrinsic excitability plasticity observed in layer IV S1_BF_ excitatory neurons is a general phenomenon of primary sensory cortices, we recorded F/I curves in layer IV excitatory pyramidal neurons in binocular primary visual cortex (V1_b_) at P12, 13, 14, 16, 19, and 20 (N = 19 mice). Here, there was a significant main effect of age (*p* = 0.029), although none of the post hoc pairwise comparisons reached significance. P14 vs P20 was the closest contrast to significance (*p* = 0.075); the mean F/I slope on P14 was lower than on P20.

Because eye opening can occur over a range of days within the window of our recordings, we aligned V1_b_ neurons based on each animal’s eye-opening status to understand if this, not age, could recapitulate the results from S1_BF_. F/I curve slopes did not differ significantly as a function of the status of eye-opening (*p* = 0.173; Fig. 5C).

Taken together, these results demonstrate a) increased excitability on P14 is not a universal feature of excitatory neurons in primary sensory cortices, b) the sudden onset of organized input into S1_BF_ does not cause plasticity of intrinsic excitability in layers II/III, and c) the sudden onset of organized visual information does not drive intrinsic excitability plasticity in excitatory layer IV neurons in V1_b_.

### Intrinsic plasticity around P14 is experience-dependent

Critical periods for postnatal isocortical plasticity are profoundly influenced by sensory experi-ence^80^. To investigate whether passive whisker input prior to P14 is required for the observed intrinsic plasticity in layer IV of the S1_BF_, we conducted unilateral whisker plucking at P10, 12, 14, 16, and 18 across a cohort of 32 mice. Subsequent F/I curves were recorded on P11, P12, P14, P16, P18, and P19, as depicted in (Fig. 6A). Importantly, whisker plucking preserves the follicles and underlying whisker musculature, maintaining the integrity of the sensory apparatus while isolating the effects of tactile input, and the contralateral whiskers are unperturbed^17^.

**Figure 6:**
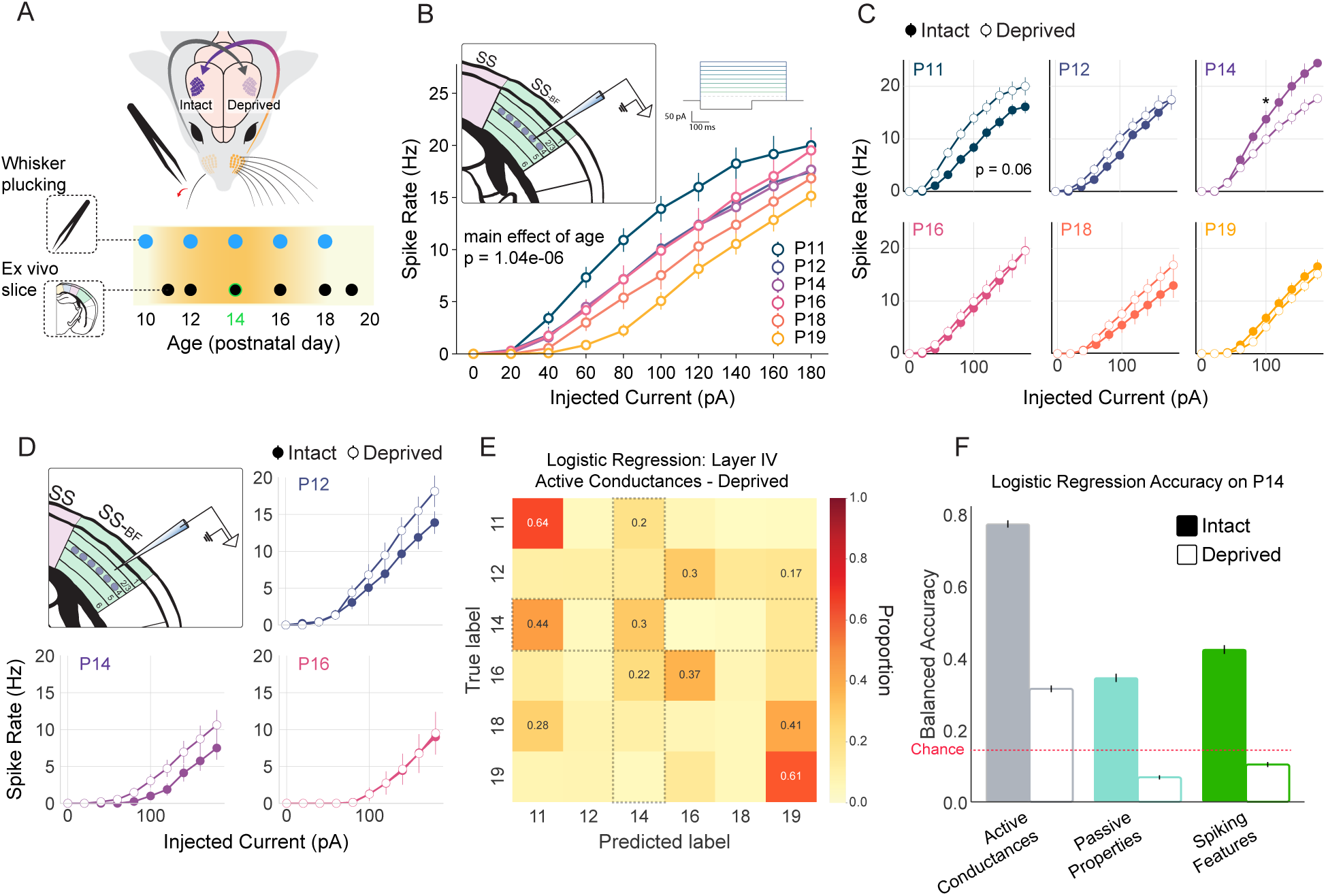
Increase in intrinsic excitability is experience dependent and is blocked by early whisker deprivation. **A)** Timeline of whisker deprivation. Mice were unilaterally whisker plucked starting at 10 days postnatal (P10) and plucking continued every other day until the day of recording. Ex vivo slices were taken from the barrel field of primary somatosensory cortex (SS_BF_) from postnatal day 11 (P11) through P19 **B)** F/I curves from Layer IV excitatory cells in the deprived hemisphere do not show the transient increase in intrinsic excitability indicating that this change is experience dependent, insets show the location of intracellular recordings, left, and current injection traces, right. **C)** Comparing F/I curves for the intact vs deprived hemisphere at each age shows a one day increase in intrinsic excitability in the deprived hemisphere on P11, top left. These comparisons also show that intrinsic excitability in the deprived hemisphere is lower than in the intact hemisphere, further suggesting this change is experience dependent. * indicates *p <* 0.05; two-sample t-test of firing frequency at 100 pA current injection. **D)** F/I curves from Layer II/III excitatory cells in the deprived hemisphere do not show an increase in excitability on P14. Intrinsic excitability in deprived hemisphere Layer II/III cells is slightly higher than in the control hemisphere, and overall excitability decreases with age. **E)** Confusion matrix showing the ability of a logistic regression classifier to predict an animal’s age based on active conductances in the deprived hemisphere (active conductances comprise dynamic input resistance and time to first spike; correct classifications fall along the diagonal). Note that chance is 16.67% and that P14 is minimally identifiable. **F)** Classifier balanced accuracy as a function of feature set (gray: active conductances, teal: passive properties, green: spiking features) and condition (intact: solid, deprived: open). Chance is indicated by the dashed red line. For Layer 4 F/I curves, N=32 mice, N=142 cells. For Layer 2/3 F/I curves, N=12 mice, N=40 cells.

In the context of sensory deprivation, layer IV S1_BF_ excitatory F/I curves revealed a significant main effect of age (*p* = 3.86*e* − 07). This comprised a lessening of intrinsic excitability across development, exemplified by the contrast of P19 (the least excitable) to P11 (the most excitable; *p* = 0.0000001; Fig. 6B). Comparing intact to deprived, there was a near-significant main effect of sensory condition (0.083), and a significant interaction of sensory condition and age (*p* = 0.006). Given the focused hypothesis on P14, a two-sample t-test was employed to compare firing rate at 100 pA between intact and deprived sensory conditions. Whisker plucking significantly reduced the excitability of layer IV barrel excitatory neurons on P14 (*p* = 0.040; Fig. 6C). In the full analysis (ANOVA), the only intact vs deprived comparison that was near significant was on P11 (*p* = 0.069). This is remarkably different than the increased excitability observed on P14 with intact whiskers (Fig. 2E).

The effect on deprived P11 in consistent with a rapid homeostatic response to offset the impact of deprivation^49,52,61^. Remarkably, despite the increase in excitability, we observed no concomitant change in mEPSC amplitude on P11 after 24 h of deprivation (p = 0.9976; Fig. S3). Taken together, these data suggest three things; first, the intrinsic plasticity on P14 requires prior experience, second, whisker deprivation drives an acute homeostatic increase in excitability, and third, there is a notable absence of synaptic homeostasis in post-critical-period layer IV S1_BF_.

We next evaluated the impact of whisker deprivation in layer II/III S1_BF_ excitatory neurons on P12, P14, and P16 in N = 12 mice. Neurons in the deprived hemisphere exhibited a significant main effect of age (*p* = 0.00124) in which intrinsic excitability declined over time, as observed under intact conditions — there was no evidence of an aberration at P14. In layer II/III, the main effect of sensory condition was also near near-significant (*p* = 0.085), and consistent with a homeostatic increase in excitability following whisker plucking. In contrast to layer IV, there was no interaction of age and sensory condition (*p* = 0.545), suggesting that the effect of whisker plucking was consistent from P12-16 in layers II/III. Across layers II/III and IV, the only condition in which intact excitability was greater than deprived excitability was on P14 in layer IV (Fig. 6C,D).

These data raise the question of how experience modifies neuronal excitability. One possibility is that the mechanisms underlying the layer IV plasticity on P14 — active conductances (Fig. 4K-M) — are suppressed in the absence of passive input beginning at P10. In this case, a classifier trained on active conductances should fail to meaningfully identify P14. Alternatively, it is possible that whisker plucking mutes the P14 effect by reducing other features, such as passive properties. In this case, P14 should still be readily identifiable by active conductances, but a corresponding, counteractive change in another feature set should also emerge. We extracted measurements of passive, active, and spiking features from F/I recordings in deprived hemispheres between P11 and P19. Trained on active conductances, age was predicted with a balanced accuracy of 34.84 ± 0.35%, and P14 was classified with 30.4 ± 0.948% accuracy (Fig. 6E). At P11, which was remarkable for its apparent homeostatic increase following 24 h of deprivation, active-conductances provided 64% classification accuracy. Passive and spiking features supported balanced accuracies of 34.31 ± 00.34% and 33.67 ± 0.37%, respectively, although in each case performance on P14 was below chance (6.83 ± 0.58% and 8.35 ± 0.64%, respectively; Fig. 6F).

Taken together, these data suggest that plasticity of active conductances on P14 requires experience, and although driven by an opposite sensory manipulation, the homeostatic response to deprivation acts through a similar set of underlying conductances.

## Discussion

Between P12 and P14, mice transition from passive to active whisker use for exploration of objects and the environment^32^. As a result, study of the barrel cortex at this age may provide insight into the interplay of primary sensory cortical circuits and the development of motor programs. Motor control of sensory organs is a fundamental component of sensation that impacts on the structure of information arriving in the isocortex. As a result, the onset of sensory organ motor control might be hypothesized to cooperate with plastic events in corresponding sensory cortices. Here, we recorded synaptic and intrinsic properties of neurons in the barrel cortical microcircuit throughout a period of postnatal development encompassing the onset of active whisking. We describe a non-synaptic form of plasticity specific to excitatory neurons of the input layer (IV) to S1_BF_, whose locus is consistent with rapid regulation of active conductances. Summarily, this population of neurons, whose inputs include excitatory afferents from VPM and POm thalamus^74,81,82^, significantly increases their input/output ratio. This effect is robustly evident on P14 but not on P12 or P16.

While the critical period for layer IV barrel structure is closed by P14^18,24,83,84^, the critical period for Hebbian modification of intracortical processing in layers II/III opens after the barrels are formed and persists through the onset of active whisking (P10-P14)^22,33,85^ (Fig. 2B). Our finding that layer IV synaptic properties are unchanged at P14 (Fig. 2F) is consistent with layer IV barrels having already exited their critical period. In other words, around P14, the anatomical structure and synaptic architecture of the barrels are established, and active whisking is poised to drive refinement of intracortical connectivity in layers II/III—a minimum of two synapses away from the thalamus^81^. This sets up an interesting problem for developing sensory circuits. External input is necessary to drive experience-dependent circuit refinement, but isocortical gain control and inhibition by ongoing activity constrains the impact of whisker-driven input in upper layers^86–89^. Speculatively, the elevated excitability of excitatory (but not inhibitory) neurons in layer IV at P14 is well positioned to briefly increase the gain of feed-forward input to layers II/III, precisely when motor-controlled whisker input is required to drive Hebbian change.

A plastic change in intrinsic properties without a corresponding shift in synaptic strength is uncommon. Intrinsic plasticity is a key feature of both homeostatic and Hebbian plasticity^45,46,49,52,90^. However, intrinsic plasticity typically co-occurs with a variety of synaptic changes including synaptic scaling, LTP, and LTD. Put simply, when intrinsic plasticity is observed, synaptic modifications also occur as part of a broader adaptive response to environmental or experimental conditions. Against this backdrop, our results suggest that these two loci of plastic change—synaptic and intrinsic—are dissociable and capable of independent regulation. We offer two plausible scenarios that could give rise to our data. First, around P14, the lack of synaptic change implies active constraint or compensation. Alternatively, the threshold for activating intrinsic plasticity might simply be lowered around P14, while developmental programs and changing external input are insufficient to drive synaptic change. Our data provide possible clues to resolve this.

Prior work demonstrates that whisker plucking/trimming drives homeostatic synaptic scaling in layers II/III^62^, layer IV^91^, and layer V^92^ of barrel cortex. However, these observations involve animals older than P28, past the closure of all critical periods in barrel cortex. In our data, whisker plucking failed to elicit a homeostatic response in synaptic strength, but prominently increased intrinsic excitability in the first 24 h, consistent with a compensatory response. Changes in intrinsic excitability are a component of the classical homeostatic response to perturbation, which normally includes synaptic compensation^49,52^. Taken with the exclusively intrinsic plasticity around P14 and prior observations of synaptic homeostasis after P28, these data suggest that synaptic modifiability is temporarily suppressed in layer IV S1_BF_ excitatory neurons when animals begin active whisking.

A comparison of the increased excitability at P14 (under intact conditions) and the compensatory increase in excitability at P11 (following 24 h of deprivation) is worthwhile. Each observation lacks measurable synaptic change. Further, electrophysiologically, it appears that active conductances are the primary mechanism by which excitability is modified in each event. However, the stimulus-response logic is inverted between the two. In the case of whisker plucking, diminished input drives a compensatory increase in excitability — this reflects the principles of maintaining a set-point in the face of perturbation^7,52^. In the case of the onset of active whisking, however, excitability increases. This should amplify input, in an apparent anti-homeostatic logic. This suggests that, in the second week of life, intrinsic properties are actively regulated by the barrel cortical circuit for both homeostatic (negative feedback) and non-homeostatic ends.

A central question that we are unable to resolve in our data is how the intrinsic plasticity on P14 is initiated. One possibility is anticipatory. In this case, increased excitability evident at P14 is the result of a developmental program whose timing requires prior input. This explanation is parallel to the timing of key plastic events in the visual system by prior experience^80^. Alternatively, the intrinsic plasticity measured at P14 is responsive. In this case, a highly plastic set of conductances react to active whisking-driven input via positive feedback, resulting in increased excitability. Future work will be required to disambiguate these explanations by plucking whiskers at P13, potentially allowing for the assembly and normal timing of an anticipatory program. Alternatively, experimental imposition of complex whisker stimulation prior to P14 would be expected to engage a reactive mechanism.

Two further limitations should be taken into consideration. First, in addition to thalamic afferents, excitatory neurons in layer IV S1_BF_ receive substantial input from other layer IV excitatory neurons^93^. As a result, the stimulus-related impact of increased intrinsic excitability in this population cannot be ascertained in ex vivo slice recordings. While our data rule out parallel changes in inhibitory neurons, the computational impact of increased excitability in the spiny stellate (excitatory) population is likely not linear as it impacts both feed-forward and local recurrent connections. And second, without preventing the intrinsic plasticity on P14, it is only possible to speculate on developmental roles. While disrupting specific developmental events can be challenging, (see, for example, visual critical periods), modeling and careful perturbations can offer empirical insight. From this perspective, it will be interesting to ask whether P14 plasticity is evident in transgenic models of neurodevelopmental disorders.

We sought to test the possibility of a interaction between the development of sensory organ motor control and corresponding primary sensory cortices. Our results suggest an unusual decoupling of synaptic and intrinsic plasticity that results in increased gain in layer IV excitatory neurons. This effect is transient, specific to cell type, layer, and region, and requires prior experience. Together, our data indicate that motor control development may influence sensory cortical development via plastic regulation of gain control.

## Acknowledgments

This work was supported by NIH BRAIN Initiative 1R01NS118442-01 (KBH), NIH/NINDS R00NS089800 (KBH), R01NS099429 (JPC), and R01MH124808 (JDD).

We thank Drs. Arianna Maffei and Steve Mennerick for their insights and conversations about this research.

## Author Contributions

MCS, KL, and KBH developed the conceptualization and methodology of the experiments. MCS, LT, HL, SC, and MEL collected the data. LT, MCS, and KBH analyzed the data. KBH, JDD, and JPC provided resources. LT, MCS, and KBH produced the figures and wrote the paper.

## Methods

### Animals

All surgical techniques and experimental procedures were conducted in accordance with protocols approved by the Washington University in St. Louis Institutional Animal Care and Use Committee, following the National Institutes of Health guidelines for the care and use of research animals. Mouse lines used in this study are: C57BL/6, C57BL/6J-Tg(Nkx2-1-cre)2Sand/J^76^, and TRAP (EGFP/RPL10A)^77^.

### Whisking Development

The emergence of whisking behavior was measured in C57/Bl6 mice between the ages of P7 and P16 in six litters (28 mice total). Video was recorded of mouse pups walking individually on a white background and during light manual restraint to observe the motion of vibrissae along with the snout. Whisking was divided into three categories based on the amplitude and coordination of whisker movements: 1) non-whisking mice only displayed whisker movements associated with respiration and head movement; 2) whisker twitching was defined by the onset of independent movement in the whisker pad but with relatively low frequencies and minimal sustained protraction; 3) active whisking consisted of rhythmic, high-amplitude cycles of protraction and retraction that can be oriented to palpate objects during exploration. An ordinal logistic regression model was applied in Python (using the statsmodel package) to determine the statistical significance of the effect of age on the development of whisking.

### Whisker Deprivation

Postnatal day 10 (P10) mice were anesthetized under transient isoflurane and whiskers on one side of the face were gently removed using forceps similar to previously described^18^. Whisker plucking is a robust manipulation that removes the entire whisker but does not cause damage to the follicles. Removal of whisker growth was repeated every other day until the day of recording.

### Slice preparation

Mice aged postnatal day 11 (P11) to P20 of either sex were anesthetized with isoflurane and decapitated. The brain was rapidly removed and submerged in ice cold, oxygenated artificial CSF (aCSF). aCSF was continuously oxygenated and contained the following (in mM): 126 NaCl, 3 KCl, 2 MgSO4, 1 NaPO4, 25 NaHCO3, 2 CaCl2 and 14 dextrose, oxygenated with 5% CO_2_/ 95% O_2_. For barrel cortex recordings, coronal brain slices (325 *µ*m) containing S1 were taken in the “across-row” plane, oriented at a 50*^◦^*angle toward coronal from the midsagittal plane. With this technique, each slice containing the posteromedial barrel subfield (PMBSF) includes one barrel column from each of the five whisker barrel rows (A through E), allowing for each column to be identified in slices and used for targeting the electrophysiological recordings^94,95^.

For visual cortex recordings, coronal brain slices (325 *µ*m) containing V1 were taken as previously described^49^. Slices were incubated at 37 *^◦^*C for 30 min and then kept at room temperature and oxygenated with 95/5 until recording (0.5-7hrs).

### Electrophysiology

Recordings were performed at 33 *^◦^*C, with constant oxygenation (95/5 O_2_/CO_2_) and circulation of aCSF. Borosilicate glass recording pipettes with tip resistances of 3-6 MΩ were pulled using a Sutter P-2000 micropipette puller. Standard internal solution contained (mM): 20 KCl, 100 K-gluconate, 10 HEPES, 4 Mg-ATP, 0.3 Na-GTP, 10 phosphocreatine, and 0.4% biocytin. Barrel cortex recordings were performed within slice regions where PMBSF structure could be clearly visualized under a 10x air objective. Neurons were visualized with a 40x immersion objective using infrared differential contrast imaging (IR-DIC). Excitatory cells in layers IV and II/III were identified based on morphology (stellate in layer IV, pyramidal in layer II/III)^26,96^. PV+ and SST+ interneurons were identified using EGFP expression driven by the NXK2.1-cre allele^97,98^. V1 was identified using the shape and morphology of ventricles and the white matter, and recordings were obtained from layer IV pyramidal neurons based on morphology^96^. F/I curves were recorded in standard aCSF containing the channel blockers APV (50 *µ*M), DNQX (20 *µ*M) and picrotoxin (20 *µ*M) in current-clamp mode. A small DC bias current was injected to maintain Vm at −70 mV between depolarizations. For excitatory F/I curves, current injections of increasing magnitude were injected from 0-180 pA with steps of 20 pA. For inhibitory F/I curves, a firing threshold was first determined using brief current injection. Current injections were then injected in 20 pA steps from threshold to 600 pA. For AMPA miniature EPSC (mEPSC) recordings, neurons were voltage clamped at −70 mV in standard aCSF containing the channel blockers APV (50 *µ*m), picrotoxin (20 *µ*M) and TTX (0.2 *µ*M). Neurons were not included in analyses if the resting membrane potential was more positive than −55 mV, input resistance was *<*80 MΩ, series resistance was *>*20 MΩ, or if any of these parameters changed *>*20% during the recording. Morphology, as well as location in the layer was confirmed by post hoc reconstruction of biocytin fills. The first 100 individual mEPSCs were isolated from recordings using the Event Search function of pClamp. All data were further analyzed using custom Python scripts.

### Histology

Recorded excitatory and inhibitory neurons were filled with biocytin (0.4% in internal solution) and slices were fixed overnight in 4% paraformaldehyde PBS. Slices were then washed in PBS, incubated in 5% NGS, 1% Triton X-100 block solution for 2 h, and incubated overnight at 4°C with AlexaFluor-568 streptavidin (1:2000, Invitrogen) and anti-somatostatin antibody (1:500, Abcam) in block solution. On the second day, slices were washed in PBS and incubated with Goat Anti-Rabbit AlexaFluor-405 (1:500, Abcam) in block solution for 1 h, 0.1% TO-PRO-3 in PBS solution for 30 minutes, then mounted on glass slides with Fluoromount and coverslipped. Slices were imaged with a fluorescent microscope (Thunder, Leica).

### Logistic Regression Classification

To predict an animal’s age based on electrophysiological features and assess the impact of sensory deprivation, we performed logistic regression analysis using Python. The analysis was conducted separately for three feature sets: active conductances (dynamic input resistance and time to first spike), passive properties (membrane resistance, membrane potential, capacitance, and tau), and spiking features (afterhyperpolarization, spike frequency adaptation, and instantaneous frequency).

Data Preprocessing: The dataset was first grouped by age and balanced by randomly sampling an equal number of cells (n) from each age group to ensure class balance. The balanced dataset was then split into training (80%) and testing (20%) sets using stratified sampling based on age. The features were standardized using the StandardScaler from the sklearn library to ensure all features had zero mean and unit variance.

Model Training and Evaluation: A multinomial logistic regression model (LogisticRegression from the sklearn library) was trained on the standardized training set. The model’s performance was evaluated on the held-out test set using 1) F1 score (macro average): The harmonic mean of precision and recall, calculated for each class and then averaged, providing an overall measure of the model’s accuracy, and 2) Balanced accuracy: The average of the proportion of correct predictions within each class, taking into account class imbalance.

The logistic regression analysis was repeated for 500 iterations (reps) to ensure robustness of the results. In each iteration, the dataset was resampled, and the model was retrained and evaluated. The mean and standard error of the metrics across iterations were reported.

Comparison of Sensory Conditions: The logistic regression analysis was performed separately for intact and deprived sensory conditions. The accuracy in predicting age 14 (the onset of active whisking) was compared between the two conditions to assess the impact of sensory deprivation on the model’s performance. The mean accuracy and standard error for age 14 were calculated across iterations.

### Statistics

Statistical analyses were performed using R and packages tidyverse (version 2.0.0) and stats (version 4.2.3).

Data Preparation: Variables ‘animal id’, ‘age’, and ‘cell id’ were converted to factors. Data were filtered to include a range of current injections in which cells were active and not saturated (e.g., between 20 and 100 pA for excitatory layer IV S1_BF_).

Slope Calculation: In the context of F/I curves, for each neuron, the slope of the frequency-current relationship was calculated using linear regression (lm function from the stats package) with ‘frequency’ as the dependent variable and ‘current’ as the independent variable.

Analysis of Variance (ANOVA): To assess the effect of age on the frequency-current slope, a mixed-effects ANOVA was performed using the aov function from the stats package. The model included ‘age’ and ‘animal id’ as fixed factors, with ‘slope’ as the dependent variable (or a neuronal property, such as capacitance where appropriate). The ANOVA model was specified as follows:

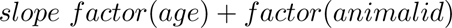

Post-hoc Tests: If the ANOVA indicated a significant main effect of, e.g., age, post-hoc tests were performed using Tukey’s Honest Significant Difference (HSD) test.

0.03 0.04 0.05 0.06 0.07

---

**Figure S1:**
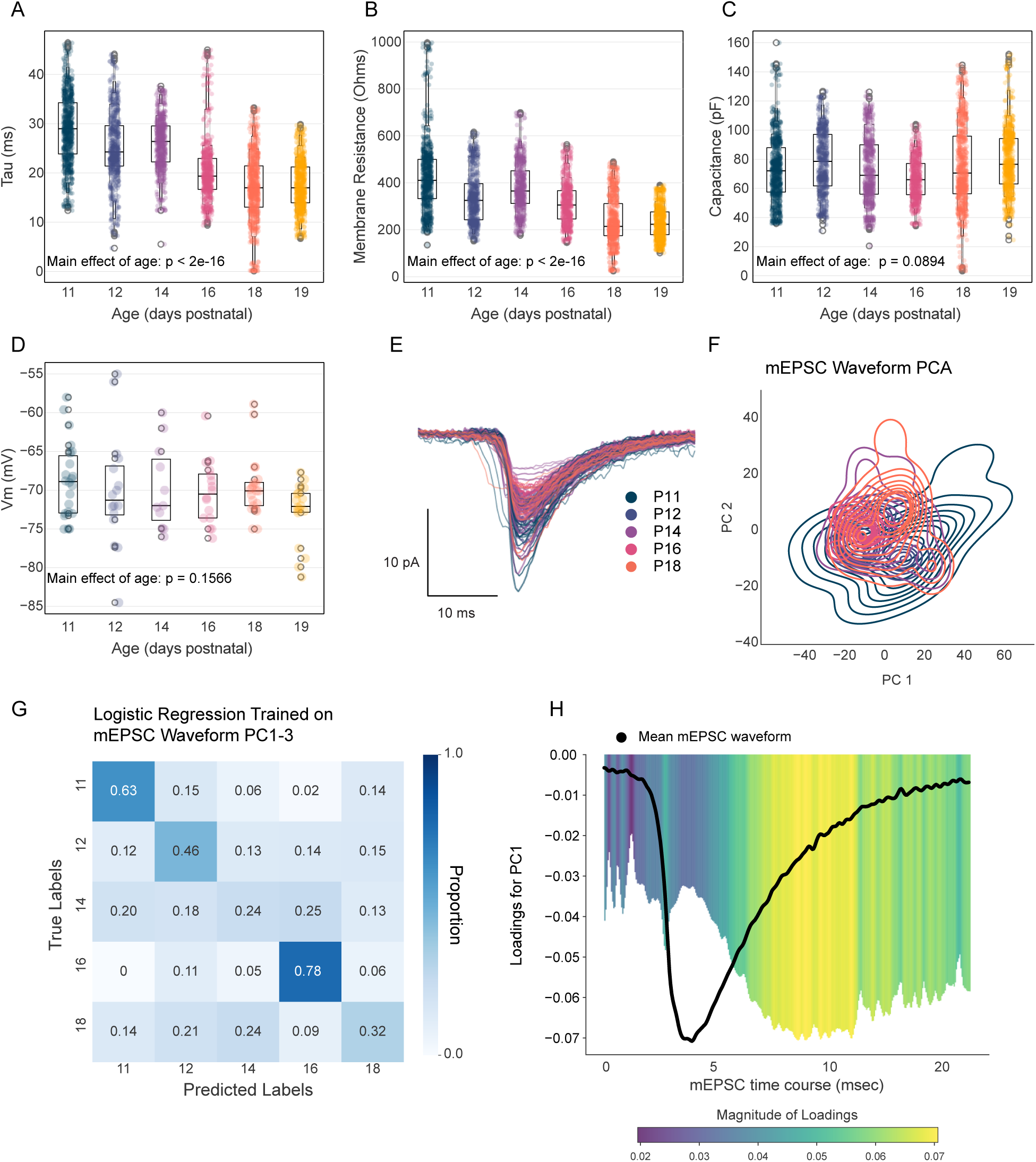
Passive membrane properties and mEPSC waveform analysis of layer IV excitatory neurons in the barrel cortex. **A-D)** Passive membrane properties of layer IV excitatory neurons as a function of age: **A)** membrane time constant (tau), **B)** input resistance, **C)** membrane capacitance, and **D)** resting membrane potential. **E)** Mean mEPSC waveforms from layer IV excitatory neurons across development (P11-P18). **F)** Kernel density estimate plot showing the the first two principal components of PCA run on the mean waveforms shown in E. Color indicates age as in E. **G)** Confusion matrix for a logistic regression model trained to predict age based on the first three principal components of mEPSC waveforms. **H)** Loadings for the first principal component (PC1), illustrating the temporal features of the mEPSC waveform that contribute most to the variance across development.

**Figure S2:**
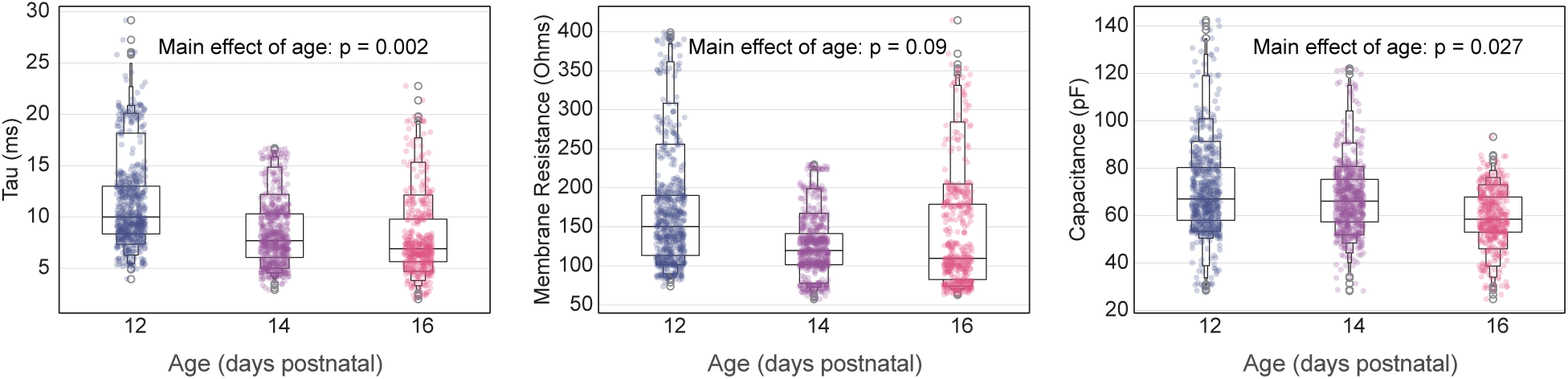
Passive membrane properties of putative parvalbumin (PV) and somatostatin (SST) interneurons in layer IV of the barrel cortex. Passive membrane properties of putative PV and SST interneurons as a function of age (P12, P14, and P16): (Left) membrane time constant (tau), (Center) input resistance, and (Right) membrane capacitance.

**Figure S3:**
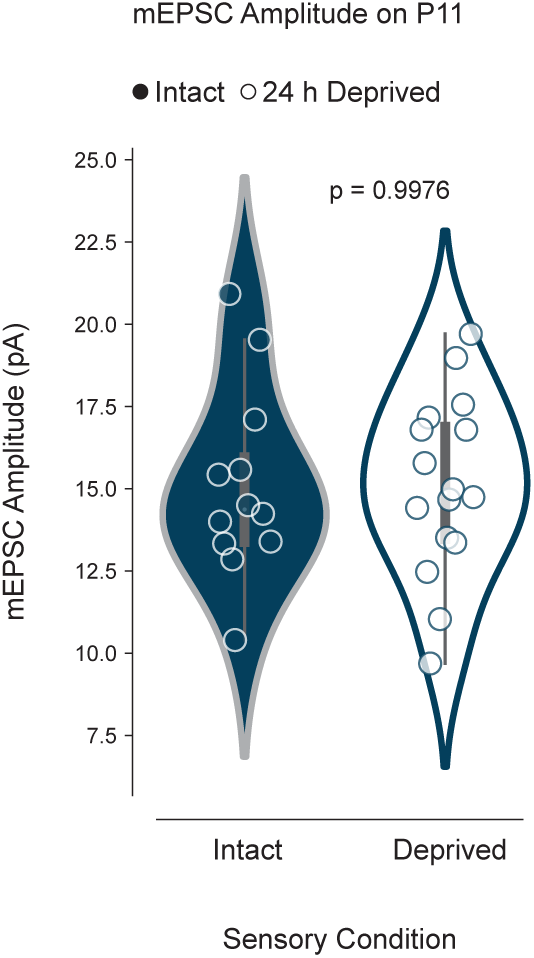
Whisker deprivation does not induce homeostatic synaptic scaling in layer IV excitatory neurons at P11. Comparison of mEPSC amplitudes between intact and deprived hemispheres in layer IV excitatory neurons at P11, 24 h after the onset of unilateral whisker deprivation. No significant difference was observed between the two conditions (p = 0.9976), indicating an absence of homeostatic synaptic scaling at this age and time point.

